# Neural dynamics of real-world object vision that guide behaviour

**DOI:** 10.1101/147298

**Authors:** Radoslaw M. Cichy, Nikolaus Kriegeskorte, Kamila M. Jozwik, Jasper J.F. van den Bosch, Ian Charest

**Author notes:** **CORRESPONDING AUTHOR** Radoslaw Martin Cichy, Department of Education and Psychology, Free University Berlin, JK 25/221b, Berlin, Germany.

## Abstract

Vision involves complex neuronal dynamics that link the sensory stream to behaviour. To capture the richness and complexity of the visual world and the behaviour it entails, we used an ecologically valid task with a rich set of real-world object images. We investigated how human brain activity, resolved in space with functional MRI and in time with magnetoencephalography, links the sensory stream to behavioural responses. We found that behaviour-related brain activity emerged rapidly in the ventral visual pathway within 200ms of stimulus onset. The link between stimuli, brain activity, and behaviour could not be accounted for by either category membership or visual features (as provided by an artificial deep neural network model). Our results identify behaviourally-relevant brain activity during object vision, and suggest that object representations guiding behaviour are complex and can neither be explained by visual features or semantic categories alone. Our findings support the view that visual representations in the ventral visual stream need to be understood in terms of their relevance to behaviour, and highlight the importance of complex behavioural assessment for human brain mapping.

## 2 Introduction

To survive, visual animals must form accurate representations of the world to guide behaviour (Heekeren et al., 2008; Panzeri et al., 2017). The underlying neural mechanisms can be formalized as a two-step mapping from sensory input to neural representations, and from neural representations to behaviour. While each of the two steps can in principle be investigated in separation, as visual recognition (DiCarlo et al., 2012) and perceptual decision making (Heekeren et al., 2008), only a combined investigation of both steps can demonstrate that experimentally measured activity during perception is in fact used by the brain to guide behaviour (de-Wit et al., 2016; Panzeri et al., 2017).

Most previous research has employed one of two strategies to investigate the triadic relationship between sensory input, neural representation and behaviour. The first approach is to relate human performance in binary classification tasks to brain activity (Newsome et al., 1989; Britten et al., 1996; Thorpe et al., 1996; Grill-Spector et al., 2000; VanRullen and Thorpe, 2001; Philiastides and Sajda, 2006; Williams et al., 2007; Ratcliff et al., 2009; Carlson et al., 2013; Ritchie et al., 2015). This strategy is experimentally elegant in its simplicity and of direct relevance to categorisation as a major function of the human visual system. However, it does not address the complexity and richness of human visual experience: our perception of the world is not binary but it is organised according to many properties, such as shape, colour and category.

A second strategy is to establish a second-order similarity between behavioural judgments and neural activity. If two stimuli with similar neural representations are judged as being similar perceptually, this suggests a role of the neural representations in guiding behaviour. This approach has established a role for the behavioural relevance of object shape representations in primate brains focusing mostly on artificial shape stimuli (Op de Beeck et al., 2001; Kayaert et al., 2005; Haushofer et al., 2008; Op de Beeck et al., 2008b p.200; Walther et al., 2009). However, it is unclear how the link between neural representations and behavior holds in a more ecologically valid setting with a large set of real-world stimuli.

Here, by combining elements of both strategies we epitomized on their respective advantages. First, to assess human behaviour related to perception of real-world visual objects, we assessed behaviour using a similarity judgments task where multiple object-pair similarities are judged at once (Kriegeskorte and Mur, 2012) for a large set of objects, establishing the link between visual input and behaviour. Second, we resolved brain responses to the same visual stimuli in space by full-brain functional magnetic resonance imaging (fMRI) and in time by magnetoencephalography (MEG), establishing the link between experimental stimuli and brain activity. Third, using representational similarity analysis (RSA; Kriegeskorte and Kievit, 2013), we forged a link between perceptual judgments and brain activity, resolving behaviour-relevant brain activity during object vision in both space and time.

## 3 Materials and Methods

### 3.1 Visual stimulus set and experimental design

The stimulus set consisted of 118 square images of everyday objects, each one from a different category, on real-world backgrounds from the ImageNet database (Fig. 1A). The same stimulus set was used in three separate experiments: behavioural, MEG and fMRI. The experiments were conducted according to the Declaration of Helsinki and approved by the respective local ethics committees (for fMRI and MEG experiments: Institutional Review Board of the Massachusetts Institute of Technology, for behavioural experiments: Ethics Committee of the Free University Berlin).

**Figure 1:**
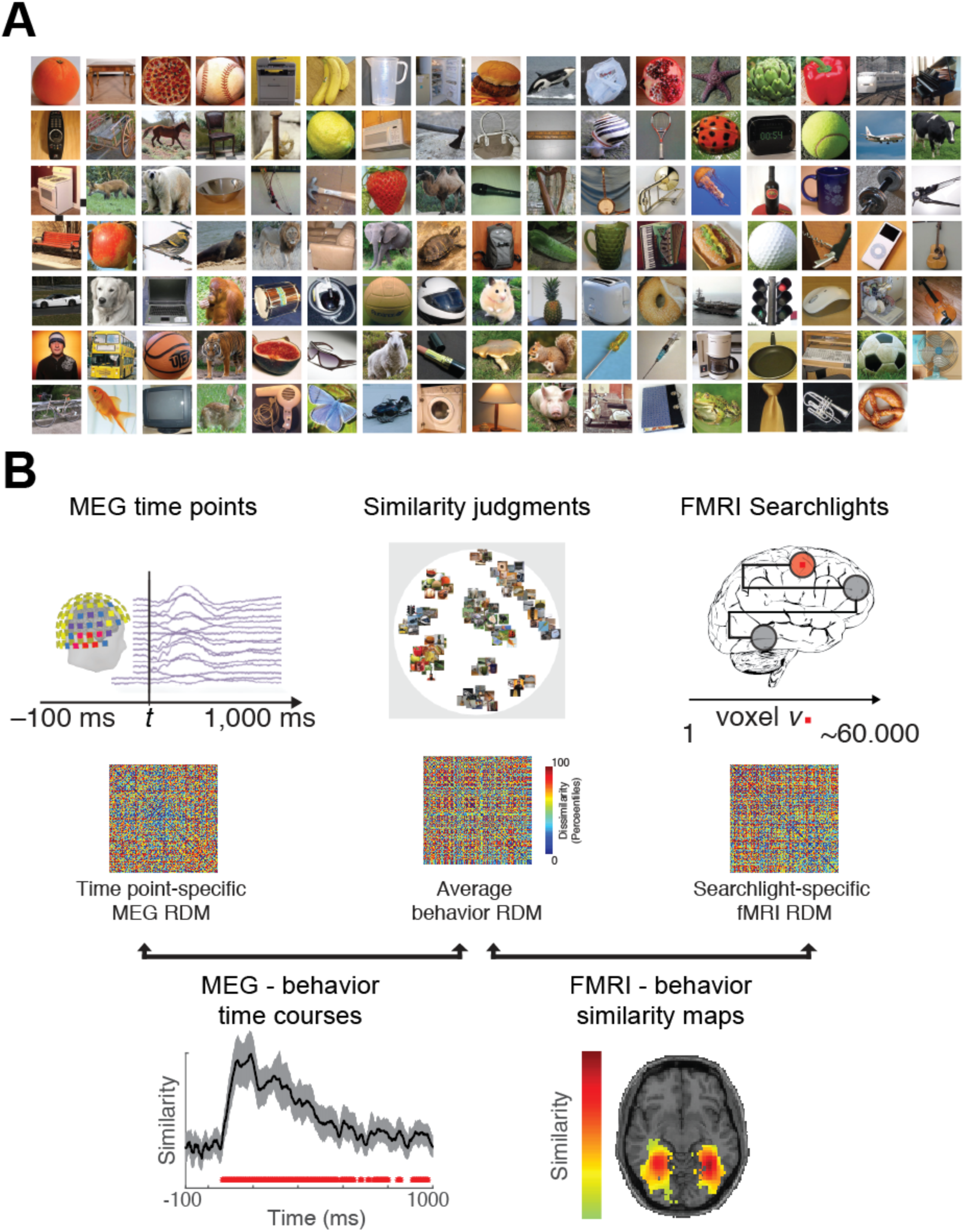
Stimulus set and analysis rationale. A) The stimulus set consisted of 118 images of real-world objects, each from a different basic level category. B) General analysis rationale. We used representational similarity analysis to link sensory input, neural activity and behaviour related to visual objects together in a common quantitative framework. We assessed behaviour using a similarity-based multiple arrangements task, and summarized the results in a behaviour RDM. To resolve the temporal aspects of behaviour-relevant brain activity, we compared the behaviour RDM with MEG RDMs in a time resolvedfashion, from - 100 to +1000ms with respect to image onset. To resolve behaviour-relevant brain activity in space, we compared the behaviour RDM with fMRI RDMs in a searchlight procedure, yielding a 3D map of representational similarity between behaviour and neural patterns. This analysis yielded time courses of representational similarity between behaviour and neural dynamics.

### 3.2 Behavioural ratings of perceived similarity

Participants (*n* = 20, 11 female, age: mean ± s.d. = 25.47 ± 7.09 years) gave ratings on the perceived similarity of object images in a multiple arrangements task (Kriegeskorte and Mur, 2012; Charest et al., 2014). Participants were asked to arrange the images on a computer screen inside a white circular arena by using computer mouse drag and drop operations. On the first trial, the participants arranged all the images at once, and subsequent trials were constructed with an adaptive selection procedure, optimizing information for all possible pairs of images. These subsequent trials often included items that were placed close together in the initial trial (sometimes in small piles). As subsequent trials included fewer objects to place, participants could refine their judgements with distinctions that are more difficult to carry in the context of the whole image set and the limited arena space. This task is efficient when dealing with large number of stimuli, and reliable similarity judgments were obtained with 60 minutes of total experiment time per participant.

### 3.3 Neuroimaging experiments: participants and experimental design

MEG and fMRI data have been published previously (Cichy et al., 2016b). Here we provide a summary of the relevant parameters.

Participants (*n* = 15, 5 female, age: mean ± s.d. = 26.6 ± 5.18 years) took part in an fMRI and MEG experiment. These participants were distinct from the participants that took part in the behavioural experiment. All participants were healthy and right handed with normal or corrected-to-normal vision. The stimulus set was identical to the one in the behavioural experiment. In both the fMRI and MEG experiments, images were presented at the center of a screen at 4.0° visual angle with 500 ms duration, overlaid with a gray fixation cross. Further presentation parameters were adjusted to the specific requirements of each imaging technique.

In the MEG experiment, each participant completed one session consisting of 15 runs of 314 s duration each. In each run, every object image was shown twice, with random condition order and a trial onset asynchrony of 0.9-1 s. Participants were instructed to respond to the image of a paper clip shown randomly every 3-5 trials (average 4) with an eye blink and a button press, and not to blink their eyes at other times.

In the fMRI experiment, each participant completed two sessions of 9-11 runs of 486 s duration each. In each run, every object image was shown once, and condition order was randomized with an inter-trial interval of 3s. In addition, 39 null trials (gray background) were interspersed randomly during which only a gray background was presented. Participants were instructed to respond to the change in luminance of the fixation cross with a button press.

### 3.4 MEG acquisition and preprocessing

MEG signals were recorded with a sampling rate of 1 kHz, filtered between 0.03 and 330 Hz, from 306 channels (204 planar gradiometers, 102 magnetometers, Elekta Neuromag TRIUX, Elekta, Stockholm). We applied temporal source space separation (maxfilter software, Elekta, Stockholm (Taulu et al., 2004; Taulu and Simola, 2006)) before analysing data with the Brainstorm software (Tadel et al., 2011). We epoched data from -100ms to +700ms with respect to the onset of each trial, removed baseline mean and smoothed data with a 20-ms sliding window. This resulted in 30 trials for each condition, session, and participant.

To determine the time-resolved similarity relations between visual representations revealed by MEG, we used multivariate pattern analysis as described previously (Cichy et al., 2014, 2016b). The rationale is that the more dissimilar two visual representations are, the more dissimilar are the resulting MEG channel activation patterns, and the better is the performance of the classifier. Thus, classifier performance can be interpreted as a dissimilarity measure of neural representations. We analysed data separately for each subject. For each time point, we extracted and arranged MEG data as 306 dimensional measurement vectors (corresponding to the 306 MEG sensors), yielding 30 raw pattern vectors per condition. We averaged raw pattern vectors by 5 before multivariate analysis to reduce computational load. We used supervised learning with a leave-one-trial-out cross-validation scheme to train and test a support vector machine (SVM) to discriminate each pair of conditions in the LibSVM implementation (http://www.csie.ntu.edu.tw/~cjlin/libsvm). Each pairwise discrimination was repeated 100 times with random assignment of raw trials to average trials, and average trials to the training and testing set. Resulting decoding accuracies were averaged across cross-validation iterations, and assigned to a matrix of size 118 *× 118*, with rows and columns indexed by the classified conditions. The matrix was symmetric and the diagonal was undefined. This procedure yielded one 118 × 118 matrix of decoding accuracies for every time-point, and we refer to it as the MEG representational dissimilarity matrix (MEG RDM).

### 3.5 fMRI acquisition & analysis

We acquired MRI data on a 3T Trio scanner (Siemens, Erlangen, Germany) with a 32-channel head coil. Structural images were acquired using a standard T1-weighted sequence (192 sagittal slices, FOV = 256 mm^2^, TR = 1,900 ms, TE = 2.52 ms, flip angle = 9°). Functional images covering the whole cortex were acquired in runs of 648 volumes using a gradient-echo EPI sequence (TR = 750 ms, TE = 30 ms, flip angle = 61°, FOV read = 192 mm, FOV phase = 100% with a partial fraction of 6/8, through-plane acceleration factor 3, bandwidth 1816Hz/Px, resolution = 3mm^3^, slice gap 20%, slices = 33, ascending acquisition).

We processed fMRI data using SPM8 <u>(http://www.fil.ion.ucl.ac.uk/spm/)</u> for each participant separately. We realigned and co-registered fMRI data to the T1 structural scan before normalizing it to the standard MNI template. To estimate condition-specific responses, we used a general linear model (GLM) into which image onsets were entered as regressors and were convolved with a hemodynamic response function. We included movement parameters as nuisance regressors. The estimated condition-specific GLM parameters were converted into *t*-values by contrasting each condition estimate against the implicitly modelled baseline.

To determine the spatial dynamics of object vision, and to relate them to behaviour, we analysed fMRI data using a spatially unbiased searchlight approach (Kriegeskorte et al., 2006; Haynes et al., 2007). In detail, for each voxel v, we extracted condition-specific *t*-value patterns in a sphere centred at v with a radius of 4 voxels (searchlight at *v*) and arranged them into pattern vectors. For each pair of conditions, we calculated the dissimilarity between pattern vectors by 1 minus Spearman’s ρ, resulting in 118 × 118 fMRI representational dissimilarity matrices (fMRI RDM) indexed in columns and rows by the compared conditions. fMRI RDMs were symmetric across the diagonal, and entries were bounded between 0 (no dissimilarity) and 2 (complete dissimilarity). This procedure resulted in one fMRI RDM for each voxel in the brain.

### 3.6 Representational similarity analysis between behaviour RDMs and neuroimaging data

To identify behaviour-relevant brain dynamics in space and in time, we related behavioural ratings to brain activity using representational similarity analysis (RSA).

To investigate temporal dynamics, we conducted time-resolved RSA between the behaviour RDM and MEG RDMs. For each subject separately, and for each time point, we correlated the time-point specific RDM with the average behaviour RDM. This resulted in a behaviour-MEG representational time course for each subject.

To investigate spatial dynamics, we conducted spatially-resolved RSA between behaviour RDM and fMRI RDMs. For this we used a searchlight analysis (Haynes and Rees, 2005a; Kriegeskorte et al., 2006). For each voxel we correlated (Spearman’s ρ) the fMRI RDM with the behaviour RDM, and saved the correlation coefficient at the position of the voxel. This resulted in a 3D representational similarity map for each participant.

### 3.7 Assessing the role of categories and visual features

To investigate the nature of the representational relations captured by RSA between neuroimaging data and behaviour, we conducted two additional analyses.

First, we investigated the level of categorical abstraction at which similarities between the brain and behaviour emerged. While the stimulus set was designed such that every object was from a different entry-level category, multiple objects fell into supra-level categories. Thus, putative representational similarities between brain and behaviour might be explained at the level of supra-categories, rather than categories. We identified the following supra-level categories in the stimulus set, guided by semantic divisions in the organisation of object knowledge in the human brain observed in neuropsychological research (Warrington and Shallice, 1984; Hart et al., 1985; Damasio, 1990; Martin et al., 1996; Caramazza and Mahon, 2003; Mahon and Caramazza, 2009). Two subjects performed category classification with the following results: animate objects (27), and inanimate objects (91), where animate are further subdivided into tools (21), food (18), music instruments (9), means of transport (9), electric appliance (11), balls (6), furniture (4) and miscellaneous (14). To assess the effect of supra-category, for each of the 9 subdivisions, we defined three model RDMs that capture the mean effects within each subdivision (e.g. for animate: within animate, within non-animate, and between animate & inanimate). Model RDMs were set to 1 for matrix elements defined by the relevant subdivision (e.g. for within animate: all matrix elements defined by animate objects, or between animate & inanimate: all matrix elements defined by comparison of animate & inanimate objects), and 0 otherwise. In total, this resulted in 27 model RDMs. We then used partial correlation analysis in the fMRI and MEG-to-behaviour RSA analysis, partialling out the effect of those 27 RDMs, and thus the effect of supra-category.

Second, we investigated whether visual features can account for the observed behaviour-brain correspondence. To characterize images by visual features, we used deep neural networks (DNN) trained on object categorisation. This model class has been shown to perform best on object categorisation tasks (Krizhevsky et al., 2012; He et al., 2015), and to predict visual object-related brain activity better than any previous model class (Khaligh-Razavi and Kriegeskorte, 2014; Yamins et al., 2014; Güçlü and Gerven, 2015; Cichy et al., 2016a). We constructed RDMs based on DNN layer-specific activation patterns of the stimulus set from an 8-layer artificial DNN trained on object categorisation on 683 object categories (for details see: (Cichy et al., 2016a), network available here: http://brainmodels.csail.mit.edu/object_dnn.tar.gz). We then used partial correlation analysis in the fMRI and MEG-to-behaviour RSA analysis, partialling out the effect of those 8 DNN RDMs.

### 3.8 Identification of behaviourally-relevant aspects of spatio-temporally resolved neural dynamics

We combined MEG and fMRI using representational similarity fusion to resolve brain dynamics related to behaviour in space and time simultaneously (Cichy et al., 2016b; Cichy and Teng, 2017). The rationale is that if signals in the MEG originate from locations resolved in fMRI, their RDMs should be similar. In addition, identifying the subset of such time point-location RDMs pairs, that also correspond to the behaviour RDMs, identifies the behaviourally relevant subset of brain activity. We performed a searchlight analysis for each fMRI subject (*n* = 15) and each time point from 0 to +500 ms in 5 ms steps. For each voxel, we correlated (Spearman’s ρ) the searchlight-specific fMRI RDM and the subject-averaged MEG RDMs. Repeated for every voxel in the brain this resulted in a 3D map of representational similarity between fMRI and MEG at a particular time point. When repeated for all time points, this resulted in a series of 3D maps revealing the spatio-temporal activation of the human brain during object perception as measured with MEG and fMRI respectively. In order to identify behaviourally-relevant aspects, we masked the results in space and time by the results of the MEG-behaviour and fMRI-behaviour RSA.

### 3.9 Statistical testing

We conducted non-parametric random effects statistics for all tests. In detail, we used right-sided sign-rank tests and corrected for multiple comparisons by FDR (p<0.05) correction. We used bootstrapping of the participant pool (10,000 iterations) to determine 95% confidence intervals on MEG-behaviour RSA peak latencies.

## 4 Results

In order to investigate the triadic relationship between sensory stimuli, neural representations and behaviour, we acquired behavioural and neural measures for a set of 118 real-world object stimuli (Fig. 1A). As a measure of behaviour, participants (*N*=20) completed a perceptual similarity judgment task. Participants arranged object images by their similarity in a 2D arena (Kriegeskorte and Mur, 2012). To assess neural activity with both high spatial and temporal resolution, we acquired full-brain fMRI and MEG measurements while participants were presented with the object images.

We related the data from MEG, fMRI and similarity judgments using RSA (Kriegeskorte, 2008; Kriegeskorte and Kievit, 2013) (Fig. 1B). In short, we computed representational dissimilarity matrices (RDMs) that capture the representational geometry between stimulus specific signals in their respective source spaces (i.e. behaviour RDMs, MEG RDMs, and fMRI RDMs). To reveal the temporal dynamics of behaviourally-relevant brain activity we compared the behaviour RDM to the time-resolved MEG RDMs (Carlson et al., 2012; Cichy et al., 2014). To identify the locus of behaviourally-relevant aspects of neural activity during object vision, we compared the behaviour RDMs to fMRI RDMs in a searchlight procedure (Haynes and Rees, 2005b; Kriegeskorte et al., 2006).

### 4.1 The temporal dynamics of behaviourally-relevant brain activity in object vision

To investigate how behaviourally-relevant brain activity during object vision emerges over time, we correlated the average behaviour RDM with MEG RDMs (N=16) from -100 to +1,000 ms in 1 ms steps with respect to stimulus onset (Fig. 1B). We found that the time course peaked at 191 ms (122 – 197 ms) (Fig. 2A), followed by a gradual decline. This demonstrates that brain activity relevant for complex behaviour emerges rapidly in the human brain, providing the basis for fast responses to the environment.

**Figure 2:**
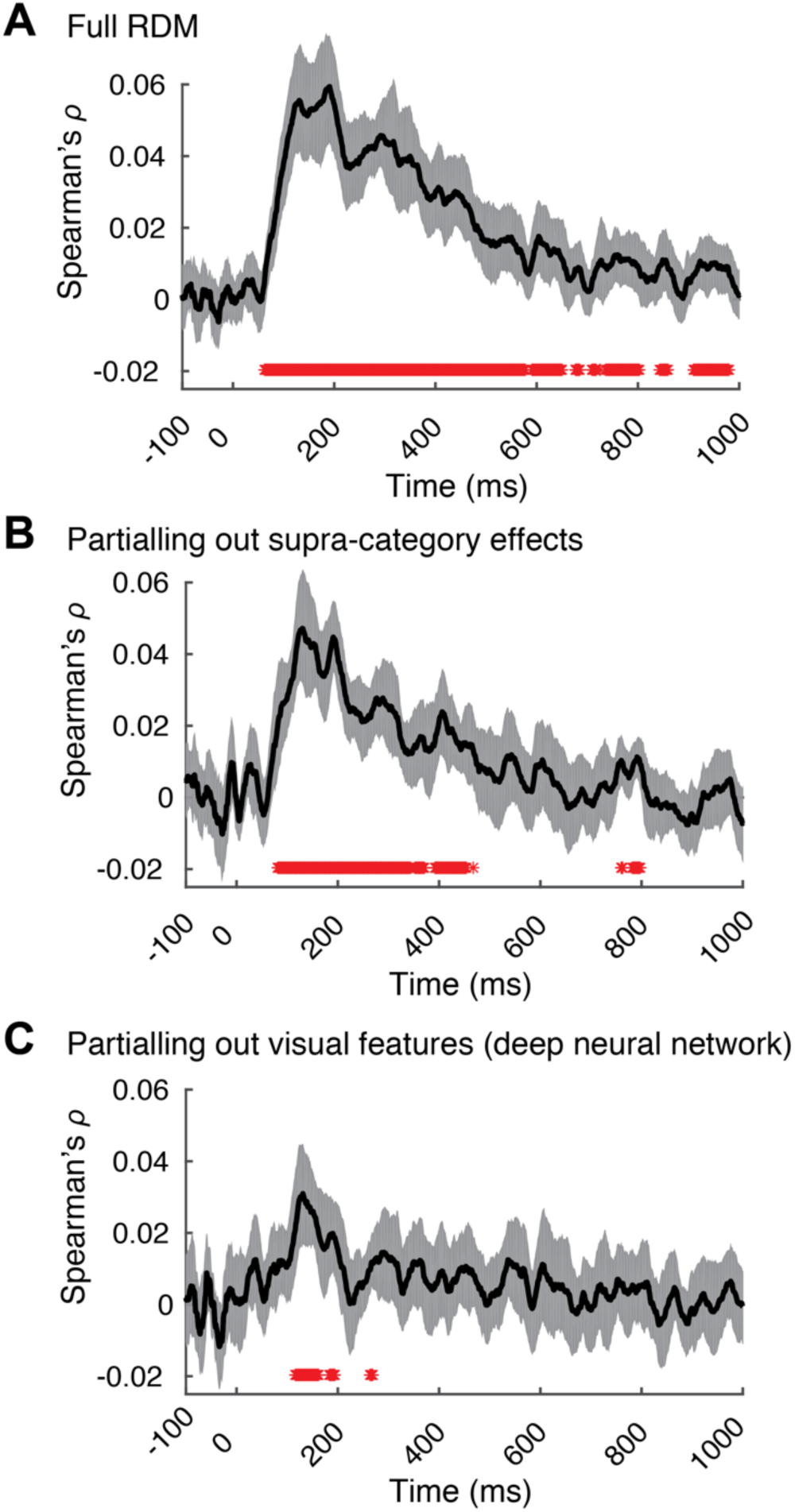
Temporal dynamics of behaviour-relevant brain activity. A) Representational similarity analysis revealed a rapid emergence of a positive correlation between perceptual similarity judgments and brain representations. A partial correlation analysis revealed that neither B) supra-category membership, nor C) visual features as assessed by a deep neural network trained on object categorization fully explained the behaviour-brain relationship.

Previous research has suggested two prominent explanations for the link between neural activity and visual behaviour. The first line of research has highlighted the role of category membership in brain and behaviour. Stimuli that belong to different categories evoke different brain responses and are judged as different (Rosch et al., 1976; Caramazza and Mahon, 2003; Grill-Spector and Malach, 2004; Op de Beeck et al., 2008a). The second line of research has put forward the importance of visual features (Tanaka, 1996; Op de Beeck et al., 2008a; Yamins et al., 2014). According to this idea, stimuli that are characterised by different visual features evoke different brain responses and are judged as different.

We performed control analyses to investigate the importance of category-membership and visual features for the emergence of the link between brain and behaviour. Concerning categories, any sizeable number of natural objects will ultimately be classified in super-ordinate groupings. Our stimulus set consisted of several supra-level categorical divisions: animals (27), tools (21), food (18), music instruments (9), means of transport (9), electric appliances (11), balls (6), furniture (4) and miscellaneous (14). To investigate the role of category-membership in the brain-behaviour relationship, we repeated our MEG-to-behaviour similarity analysis while partialling out the effect of putative differences due to supra-level category membership. This analysis revealed a significant effect with a peak at 130 ms (122 – 196 ms) (Fig. 2B), indicating that supra-category membership is not the only factor driving the brain-behaviour relationship.

To investigate the role of visual features in the brain-behaviour relationship, we used an artificial deep neural network (DNN) trained to classify objects as a proxy for these objects’ visual features. DNNs have been shown to explain visual representations in the brain better than any other previous models (Khaligh-Razavi and Kriegeskorte, 2014; Yamins et al., 2014; Güçlü and Gerven, 2015; Cichy et al., 2016a), and to predict human behaviour considerably well (Khosla et al., 2015; Kubilius et al., 2016; Peterson et al., 2016). To investigate the role of visual features, we repeated the MEG-to-behaviour analysis while partialling out the effect of DNN visual features. This revealed a significant effect with a peak at 131 ms (123 – 568 ms) (Fig 3C), demonstrating that the link between brain activity and behaviour cannot be fully explained by visual features from DNNs.

**Figure 3:**
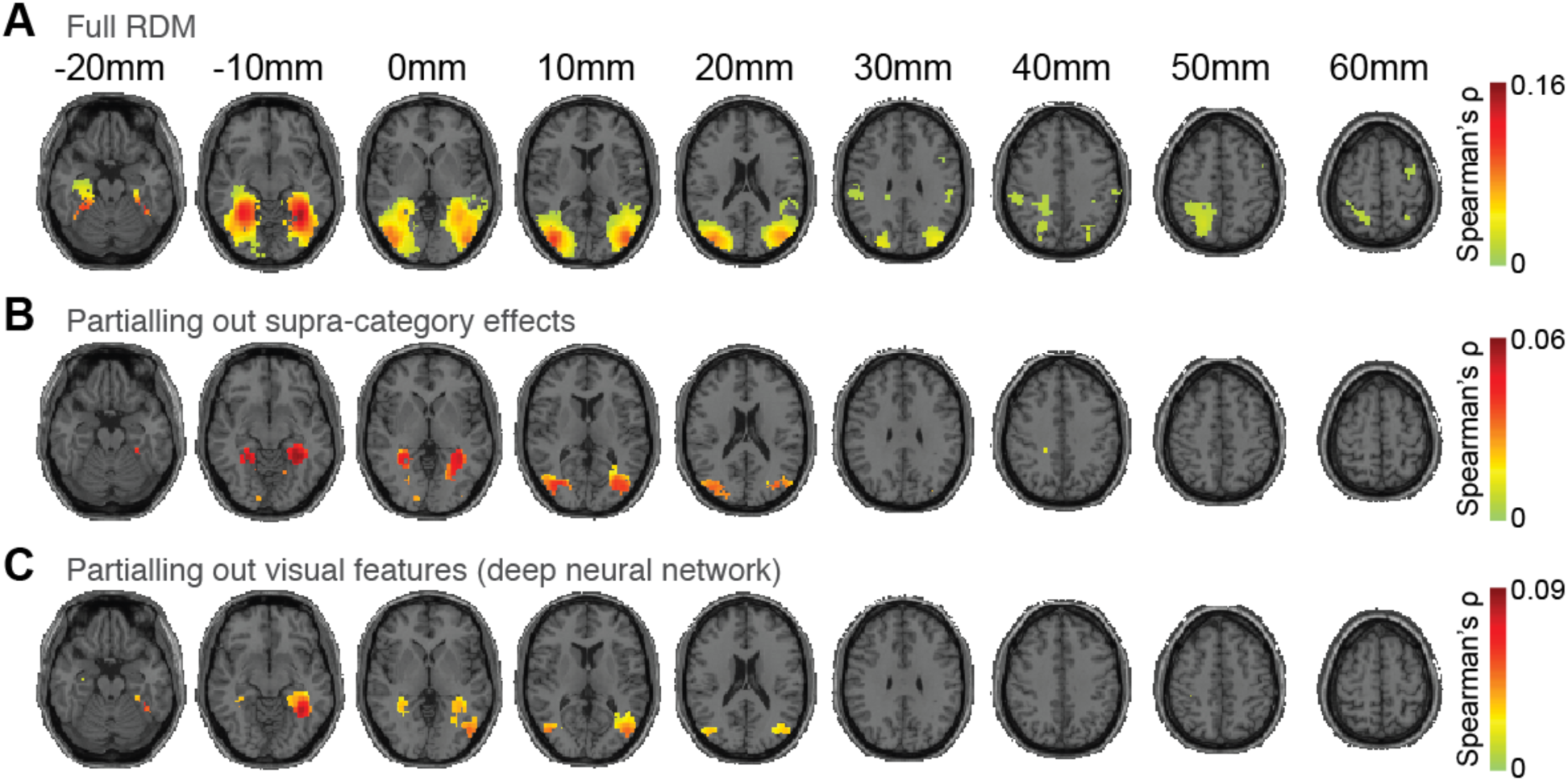
Spatial dynamics of behaviour-relevant brain activity. A) Searchlight-based representational similarity analysis revealed a positive correlation between perceptual similarity judgments and brain representations with highest correlation in high-level ventral visual cortex. A partial correlation analysis revealed that neither B) supra-category membership, nor C) visual features discovered by a deep neural network trained on object categorisation fully explained the behaviour-brain relationship.

**Figure 4:**
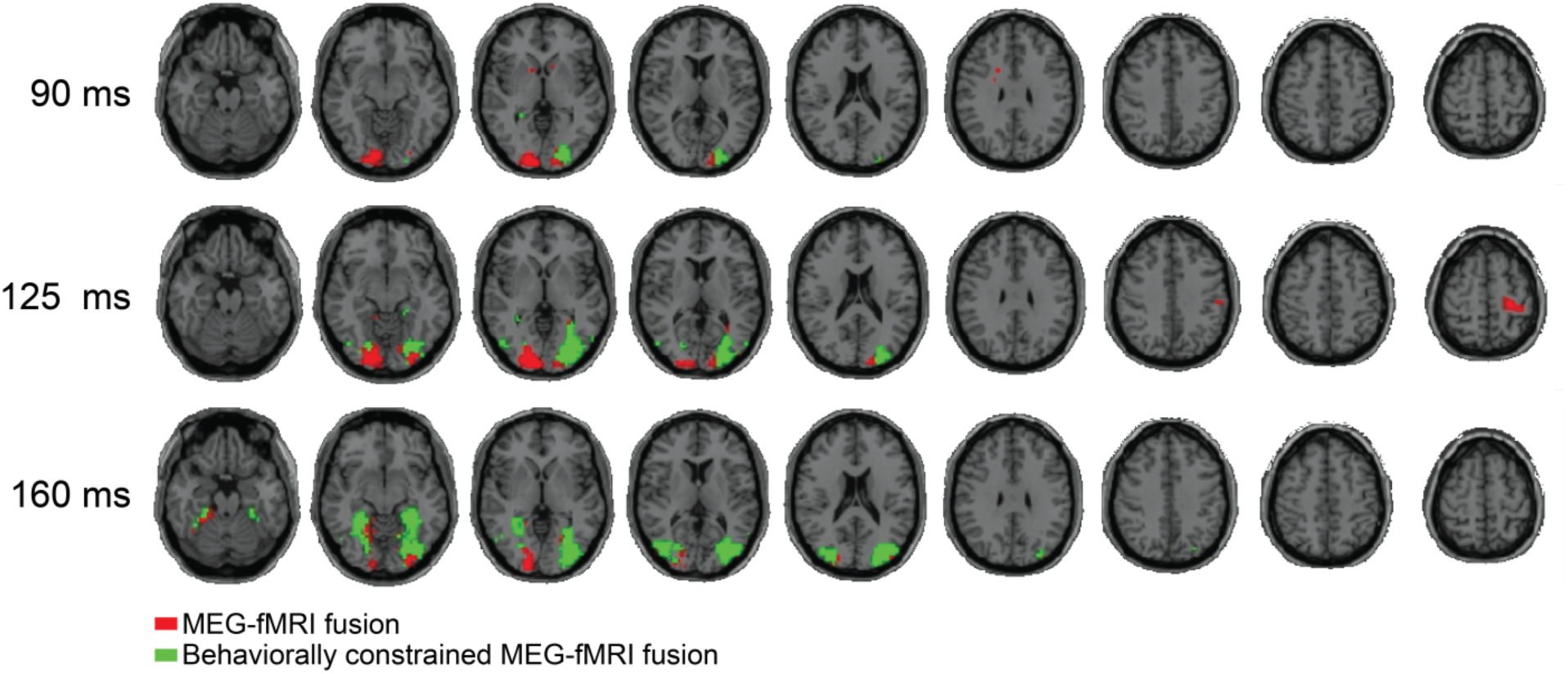
Spatio-temporally resolved behaviour-relevant brain activity during object vision. We used representational-similarity-based MEG-fMRIfusion (N = 16, p<0.05, FDR-corrected) to identify the spatio-temporal dynamics of brain activity during object vision. Red voxels indicate the significant spatio-temporal maps reflecting MEG-fMRI correspondence. Green voxels indicate the behaviourally-relevant subset of those maps, identified by masking the results of the MEG-fMRI fusion with time points and locations identified as behaviourally relevant in the analyses relating MEG and fMRI to behavioural similarity ratings separately. For a fully temporally resolved view see Movie 1.

Together, these results indicate that the emergence of behaviourally-relevant aspects of object representations is rapid, and neither fully explained by supra-ordinate categorisation, nor by visual features.

### 4.2 The spatial dynamics of behaviourally-relevant brain activity in object visions

We then investigated the role of category membership and visual features in the spatially resolved brain-behaviour relationship. Neither partialling out the effects of supra-level category membership (Fig. 3B), nor the visual features (Fig. 3C) abolished the significant brain-to-behaviour relationship in high-level ventral visual cortex.

Together, these results link behaviour to object representations in high level visual cortex and show that neither category membership nor visual feature similarity can fully account for this link.

### 4.3 Integrating spatial and temporal aspects of behaviour-relevant brain activity

The aforementioned analyses revealed that behaviour-relevant brain activity was present over a long period of time and in several brain regions. These findings raise the question of how brain regions map onto temporal processing stages: at which time point is which region of the brain the source of behaviour-relevant brain activity? To resolve the many-to-many mapping between MEG time points and brain regions, we used RSA based MEG-fMRI fusion (Cichy et al., 2014, 2016b). This approach allows identifying the spatio-temporal components of brain activity during object vision that guide behaviour (Philiastides and Sajda, 2007; Cichy et al., 2016b). First, we computed the representational similarity between fMRI locations and MEG time points (*N* = 15, right-sided signed-rank test, *p* < 0.05 FDR corrected), providing spatio-temporal maps of neural dynamics during object vision (Fig. 3A, red-colored voxels, shown for 90, 125 and 160ms; for full time-resolved results see Movie 1). Second, we masked the MEG-fMRI fusion results by the temporal and spatial relationships identified in our MEG-behaviour and fMRI-behaviour analyses respectively. This masking isolates the behaviourally relevant components in the spatio-temporal dynamics of object recognition (Fig. 3A, green-coloured voxels). We found that a subset of the brain activity is behaviourally relevant, spreading rapidly from the occipital pole in anterior direction to high level ventral visual cortex.

## 5 Discussion

Resolving how the brain translates the sensory stream hitting the retina into behaviour requires the establishment of quantitative relationships between sensory input, neural activity, and behaviour (Ince et al., 2015; de-Wit et al., 2016; Panzeri et al., 2017). Here, we established this link for ecologically valid visual input and behaviour with neural activity in both space and time.

We suggest the identified behavioural relevant neural activity may play a role in a diverse set of task contexts, as participants performed two different tasks during neuroimaging experiments and behavioural assessments. For behaviour, participants performed a multi-arrangement task juding the perceptual similarity of objects. In MEG and fMRI, participants performed a target detection task which does not involve any active similarity judgments. Because behavioural relevance was established across these very different task contexts, this strongly suggests that the identified representations are behavioural relevant in other tasks as well (Bracci et al., 2017; Kay and Yeatman, 2017). Furthermore, the identified neural activity likely generalizes across subjects, as participants differed across experiments. Thus, the identified neural representations are general and may guide complex human behaviour. Previous studies have identified behaviourally relevant spatiotemporal dynamics of primate brain activity during vision for binary classification tasks (Newsome et al., 1989; Britten et al., 1996; Thorpe et al., 1996; Grill-Spector et al., 2000; VanRullen and Thorpe, 2001; Philiastides and Sajda, 2006; Williams et al., 2007; Ratcliff et al., 2009; Carlson et al., 2013; Ritchie et al., 2015); and for similarity ratings (Op de Beeck et al., 2001; Kayaert et al., 2005; Haushofer et al., 2008; Op de Beeck et al., 2008b p.200; Walther et al., 2009; Mur et al., 2013). Here, we extend those studies in ecological validity by simultaneously using complex arrangement tasks to probe perceived similarity of real-world stimuli, increasing the ecological validity.

What are the properties of the objects according to which the link between neural activity and behaviour can be established? Two major factors that govern both behaviour and neural activity in high-level ventral visual cortex are category membership (Rosch et al., 1976; Caramazza and Mahon, 2003; Grill-Spector and Malach, 2004; Op de Beeck et al., 2008a) and visual feature similarity (Tanaka, 1996; Op de Beeck et al., 2008a; Yamins et al., 2014). By using a stimulus set where each stimulus was from a different category and by controlling for the effects of supra-ordinate category, we have shown that the brain-behaviour relationship does not solely depend on category membership. This is consistent with the result that neurons in IT carry information about a large set of object properties other than category membership alone (Op de Beeck et al., 2008a; Hong et al., 2016), and suggests that categorisation is only one of many functions of the ventral visual stream.

To address feature similarity, we investigated whether visual features from a deep neural network (DNN) trained on object categorisation could account for the brain-behavioural link. The DNN did not fully account for this link, which is consistent with the observation that such models do not fully explain activity in the primate visual pathway (Khaligh-Razavi and Kriegeskorte, 2014; Yamins et al., 2014; Güçlü and Gerven, 2015; Cichy et al., 2016a). This might be so because the visual features emerging in the DNN are optimised for the task that the network is trained on, i.e. object categorisation, while categorisation may be only one of the functions of the ventral visual pathway. The fit between brain activity and artificial neural networks might be improved by training DNNs on tasks with higher ecological validity that better capture the richness of human perception than categorisation, such as perceptual similarity arrangements.

In sum, resolving the neural activity that links sensory experience to behaviour requires differentiating behaviourally relevant neural activity from epiphenomenal activity. Here, by combining brain activity measurements from multiple imaging modalities with an ecologically valid behavioural assessment, we resolved the spatiotemporal dynamics of behaviourally-relevant brain activity for object vision. Our results further strengthen the link between brain activity in ventral visual cortex during object vision and behaviour, and highlight the importance of complex behavioural assessment for human brain mapping.

## 6 Acknowledgments

This work was funded by the German Research Foundation (DFG, CI241/1-1) to R.M.C. and by the European Research Council (ERC, 680906) to I.C and N.K. MEG and fMRI data were collected at the Athinoula A. Martinos Imaging Center at the McGovern Institute for Brain Research, MIT.

